# A Retrotranslocation Assay that Predicts Defective VCP/p97-Mediated Trafficking of a Retroviral Signal Peptide

**DOI:** 10.1101/2021.11.05.467538

**Authors:** Poulami Das, Wendy Kaichun Xu, Amit Kumar Singh Gautam, Mary M. Lozano, Jaquelin P. Dudley

## Abstract

Studies of viral replication have provided critical insights into host processes, including protein trafficking and turnover. Mouse mammary tumor virus (MMTV) is a betaretrovirus that encodes a functional 98-amino acid signal peptide (SP). MMTV SP is generated from both Rem and envelope precursor proteins by signal peptidase cleavage in the endoplasmic reticulum (ER) membrane. We previously showed that SP functions as an HIV-1 Rev-like protein that is dependent on the AAA ATPase VCP/p97 to subvert ER-associated degradation (ERAD). SP contains a nuclear/nucleolar localization sequence (NLS/NoLS) within the N-terminal 45 amino acids. To directly determine the SP regions needed for membrane extraction and trafficking, we developed a quantitative retrotranslocation assay with biotin acceptor peptide (BAP)-tagged SP proteins. Use of alanine substitution mutants of BAP-tagged MMTV SP in retrotranslocation assays revealed that mutation of amino acids 57 and 58 (M57-58) interfered with ER membrane extraction, whereas adjacent mutations did not. The M57-58 mutant also showed reduced interaction with VCP/p97 in co-immunoprecipitation experiments. Using transfection and reporter assays to measure activity of BAP-tagged proteins, both M57-58 and an adjacent mutant (M59-61) were functionally defective compared to wild-type SP. Confocal microscopy revealed defects in SP nuclear trafficking and abnormal localization of both M57-58 and M59-61. Furthermore, purified GST-tagged M57-58 and M59-61 demonstrated reduced ability to oligomerize compared to tagged wild-type SP. These experiments suggest that SP amino acids 57-58 are critical for VCP/p97 interaction and retrotranslocation, whereas residues 57-61 are critical for oligomerization and nuclear trafficking independent of the NLS/NoLS. Our results emphasize the complex host interactions with long signal peptides.

**IMPORTANCE:** Endoplasmic reticulum-associated degradation (ERAD) is a form of cellular protein quality control that is manipulated by viruses, including the betaretrovirus, mouse mammary tumor virus (MMTV). MMTV-encoded signal peptide (SP) has been shown to interact with an essential ERAD factor, VCP/p97 ATPase, to mediate its extraction from the ER membrane, also known as retrotranslocation, for RNA-binding and nuclear function. In this manuscript, we developed a quantitative retrotranslocation assay that identified an SP substitution mutant, which is defective for VCP interaction as well as nuclear trafficking, oligomer formation, and function. An adjacent SP mutant was competent for retrotranslocation and VCP interaction, but shared the other defects. Our results revealed the requirement for VCP during SP trafficking and the complex cellular pathways used by long signal peptides.

## INTRODUCTION

Mouse mammary tumor virus (MMTV) is a complex murine betaretrovirus that encodes both regulatory and accessory proteins for efficient replication and transmission in mice (1–5). Rem is a precursor protein that is translated from a doubly spliced version of envelope (*env*) mRNA on the ER membrane (1, 2). Rem then is cleaved by signal peptidase into an N-terminal signal peptide (SP) (6, 7) and a C-terminal product (Rem-CT) (4, 5). Since Rem and MMTV envelope protein are made from the same open reading frame, SP is synthesized from both singly spliced *env* mRNA and doubly spliced *rem* mRNA (2). The cleaved Env protein serves as the viral anti-receptor for mouse transferrin receptor 1 (8), whereas failure to express Rem C-terminal sequences leads to hypermutation of the MMTV genome by the Apobec family member, activation-induced cytidine deaminase (AID) after infection of BALB/c mice (4). Rem-CT is primarily located in the ER, but traffics to early and late endosomes for an unknown function (5). Our results suggest that Rem has a human immunodeficiency virus type 1 (HIV-1) Vif-like accessory function (9–11). Rem-CT also has an accessory function that is not required for MMTV replication (4, 5). MMTV SP expression is necessary for viral replication in tissue culture (1, 2, 6), whereas Rem C-terminal sequences are needed for efficient viral transmission *in vivo* (4, 12).

Conventional signal peptides are short, 20-30 amino-acid sequences that are bound by the cytosolic signal recognition particle to allow docking and co-translational transfer of ER membrane-spanning or secreted polypeptides (13, 14). Typically signal peptides are cleaved by ER-luminal signal peptidase (15) and then degraded by intramembrane signal peptide peptidases (16). MMTV-encoded SP is 98 amino acids in length and has an arginine-rich motif (ARM), typical of RNA-binding proteins, as well as a nuclear/nucleolar localization sequence (NLS/NoLS) and a leucine-rich nuclear export signal (NES) (Fig. 1A) (1, 17). These features are typical of viral nuclear export proteins, such as HIV-1 Rev (18–20). Our previous experiments have shown that SP levels are highest in the nucleolus (1). Analysis of mutants combined with transfection and reporter assays revealed that SP trafficking and function require an intact NLS and cleavage by signal peptidase (1, 2). We also have demonstrated that uncleaved Rem accumulates in the presence of proteasome inhibitors and that a dominant-negative form of the AAA ATPase family member, VCP/p97, inhibits SP activity (2, 21). These characteristics suggested that Rem is a target of the cellular protein quality control system, ER-associated degradation (ERAD) (22, 23). However, after cleavage from the precursor in the ER membrane, SP is stable and avoids proteasomal degradation (2, 21, 23). Knockdown and knockout methods in tissue culture indicate that SP activity does not need Derlin proteins (21) as observed for most ERAD substrates (24). Our experiments indicate that only Rem, not SP, is ubiquitylated, but both require the p97/VCP ATPase for activity (2, 21).

**FIG 1.**
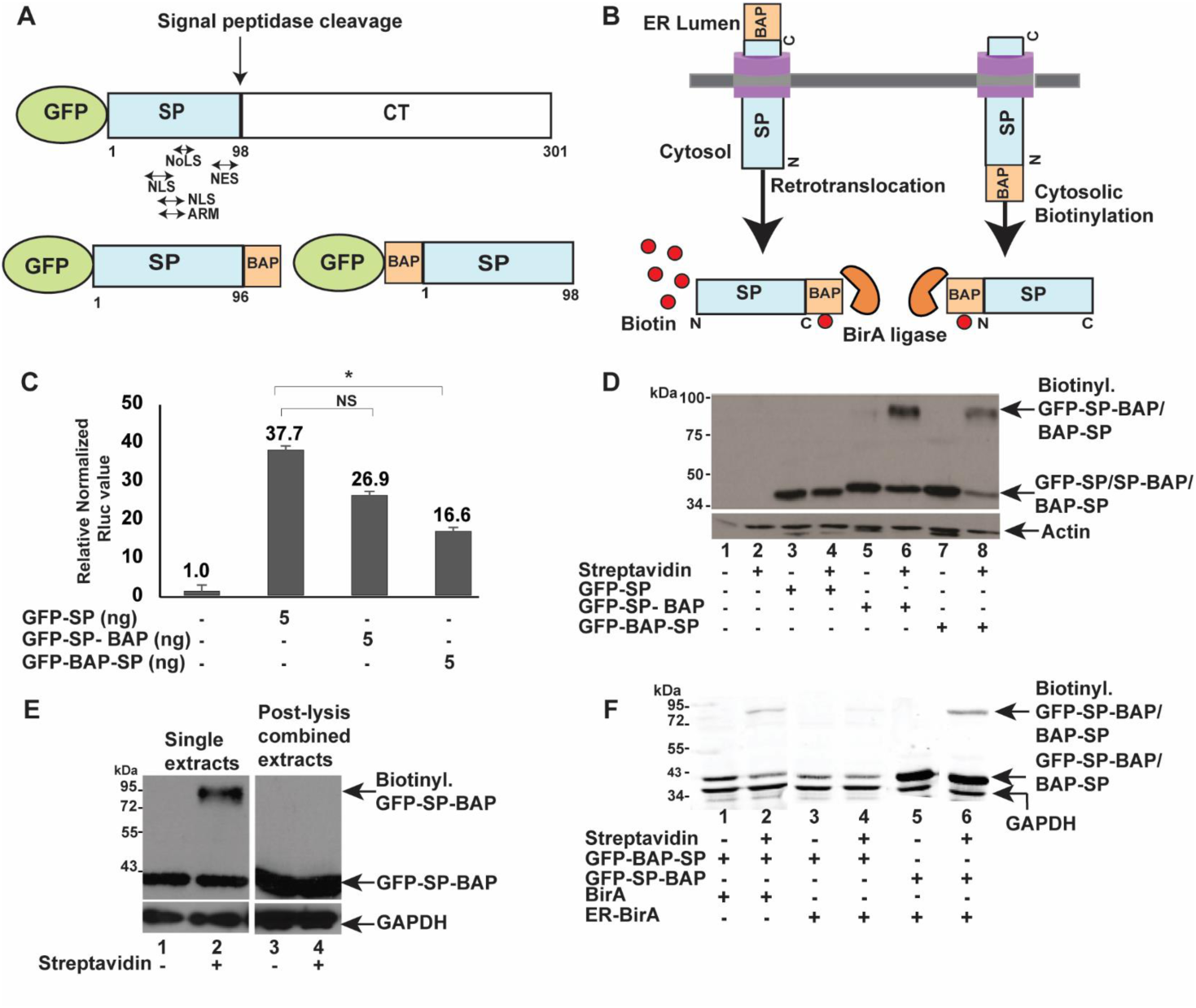
Biotinylation assay to detect MMTV SP biotinylation inside cells. (A) Diagram of GFP-tagged Rem and SP proteins. The GFP-tagged Rem precursor is shown with an arrow indicating the position of host signal peptidase cleavage between SP and the Rem C-terminus (CT). The relative positions of the nucleolar/nuclear localization sequences (NoLS/NLS), arginine-rich motif (ARM), and nuclear export sequence (NES) within SP are indicated. A BAP tag was inserted after amino acid 96 at the C-terminus of GFP-tagged SP for expression of GFP-SP-BAP. Alternatively, BAP was inserted at the N-terminus of SP to allow expression of GFP-BAP-SP. (B) Rationale for SP retrotranslocation assays. Expression vectors for GFP-SP-BAP and a cytosolic biotin-ligase (BirA) (orange Pac-Man) were co-transfected into HEK 293 cells. Following retrotranslocation, BirA covalently adds biotin (red circles) to the single acceptor lysine within the BAP tag of GFP-SP-BAP. The purple transmembrane shapes represent retrotranslocons. Co-transfection of expression vectors for GFP-BAP-SP and BirA do not require retrotranslocation for detection of biotinylated SP. Biotinylation is detected by an increased mass after incubation with streptavidin. (C) Luciferase assays confirm the activity of BAP-tagged SP proteins. Each bar represents the mean of triplicate transfections ± standard deviations in HEK 293 cells. The *Renilla* luciferase values are normalized to values for co-transfected firefly luciferase vector lacking a Rem-responsive element and in the absence of SP expression (assigned a relative value of 1). The amounts of expression plasmids used for transfection are shown in nanograms (ng). NS = not significant. * = p-value <0.05. (D) Western blots confirm SP retrotranslocation. Addition of streptavidin to extracts increases the molecular mass of GFP-SP-BAP by ∼50 kDa. Extracts from HEK 293 cells transfected with pcDNA3 (empty vector) (lanes 1 and 2) or a GFP-SP expression plasmid lacking a BAP tag (lanes 3 and 4) and BirA expression vector were used as negative controls. The upper and lower blots were incubated with antibodies specific for GFP and actin, respectively. The biotinylated and non-biotinylated forms of N-terminally and C-terminally BAP-tagged proteins are indicated by arrows. Non-biotinylated GFP-SP is detectable since biotin was added for the last 12 h of a 48 h transfection period. (E) Biotinylation of GFP-SP-BAP occurs within co-transfected cells and not after cell lysis. Lysates from co-transfections of expression plasmids for GFP-SP-BAP and BirA (lanes 1 and 2) or separate transfections (lanes 3 and 4) were prepared. Lysates from separate transfections were combined post-lysis prior to streptavidin addition as indicated. Western blots were incubated with antibodies for GFP or GAPDH. (F) ER-localized BirA biotinylates C-terminally, but not N-terminally BAP-tagged SP. HEK 293T cells were co-transfected with expression vectors for GFP-BAP-SP or GFP-SP-BAP and either BirA (cytosolic) or ER-BirA (ER localized) as indicated. Streptavidin was added to lysates in even lanes, and samples were analyzed by Western blotting using GFP- and GAPDH-specific antibodies and LI-COR detection.

VCP/p97/Cdc48 ATPase is well conserved in eukaryotes from yeast to humans (25, 26) and is used for multiple cellular functions, including ERAD (22, 22). VCP mutations are associated with human diseases, such as multisystem disease inclusion body myopathy, Paget’s disease of bone, and frontotemporal dementia (IBMPFD), amyotropic lateral sclerosis, and Parkinson’s disease (27–29). The p97 ATPase provides the energy required for the energetically unfavorable retrotranslocation step of the ERAD pathway (25). Ubiquitylated substrates are marked for degradation by specific E3 ligases within the ER membrane and then are bound to p97 through a variety of mediators, including the heterodimer UFD1–NPL4 and UBX2 (30, 31). Retrotranslocation of ubiquitylated substrates by p97/VCP to the cytosol leads to their interaction with the proteasome and degradation in the cytosol. Nevertheless, retrotranslocated SP survives p97-assisted ER membrane extraction relative to ERAD substrates (2, 3). Further, our results from a UBAIT interaction screen in human cells suggested that MMTV SP binds p97/VCP, but other ERAD-associated proteins were not identified (21).

In this report, we further explored our hypothesis that the ATPase p97 interacts with MMTV-encoded SP to mediate its retrotranslocation into the cytosol. To detect and quantify the fraction of retrotranslocated SP, we developed an assay based on specific biotin labeling of SP bearing a biotin acceptor peptide (BAP) tag on the C-terminus (SP-BAP) by the cytosolic enzyme BirA (32). Using the retrotranslocation assay and cellular extracts expressing SP-BAP mutants, we identified a 2-amino-acid region of SP that is critical for retrotranslocation and function in reporter assays. We also showed that wild-type SP interacts with p97/VCP using co-immunoprecipitation experiments and that the mutant with compromised retrotranslocation has defective interactions with p97. Reporter assays, confocal microscopy, and oligomerization assays suggested amino acids 57-61 are important for nuclear trafficking. These experiments support the idea that MMTV-encoded SP avoids proteasomal degradation by direct or indirect binding to the VCP/p97 ATPase for retrotranslocation and function of a long signal peptide.

## RESULTS

### MMTV SP retrotranslocation is detectable using a biotinylation assay

Our published experiments indicate that SP is synthesized as a type II transmembrane protein on ER membranes as part of the Env or Rem precursors and retrotranslocated to the cytosol using the p97 ATPase (2, 21, 23). These conclusions were based on the observed glycosylation of the Rem C-terminus, the ability of mutations in the predicted signal peptidase cleavage site between SP and Rem-CT to abolish SP activity, and decreased SP-reporter activity in the presence of dominant-negative VCP/p97 expression (2, 21). The SP reporter assay depends on transfection of the pHM*Rluc* plasmid, which has the *Renilla* luciferase gene inserted between splice acceptor and donor sites within the 3’ end of the MMTV genome containing the Rem-responsive element (RmRE) (1, 6). Co-expression of MMTV SP in the presence of pHM*Rluc* leads to luciferase expression if the unspliced *Renilla*-containing mRNA is exported from the nucleus for translation (1). Although highly sensitive, quantitative, and specific, this method provides an indirect measure of SP retrotranslocation.

To develop a quantitative SP retrotranslocation assay, we used an approach similar to that adopted by Petris et al. for four different cellular proteins (33, 34). A BAP tag (35) was inserted after amino acid 96 of the N-terminally GFP-tagged SP sequence (GFP-SP-BAP) using site-directed mutagenesis (Fig.1A). The placement of the tag was designed to delete the Rem signal peptidase cleavage site that would remove the BAP tag during cotranslational transfer across the ER membrane (2). We predicted that co-expression of GFP-SP-BAP with *E. coli* biotin ligase (BirA) (32) in the presence of biotin would allow BirA-mediated biotinylation of the single acceptor lysine within the C-terminal BAP tag after SP retrotranslocation to the cytosol. Biotinylation increases the mass of proteins after incubation of cellular lysates with streptavidin, which has a high affinity and specificity for biotin (36). Since the BirA ligase and the BAP tag are initially in different cellular compartments (cytosol and ER lumen, respectively), we expected that biotinylation of GFP-SP-BAP would confirm that SP is retrotranslocated (Fig.1B, left).

Our previously published experiments indicated that both N-terminally GFP-tagged Rem or SP alone are functional in the pHM*Rluc* reporter assay (2, 21). To determine whether addition of the C-terminal BAP tag interferes with SP activity, we co-transfected human embryonic kidney (HEK) 293 cells in triplicate with GFP-SP or GFP-SP-BAP expression plasmids in the presence of the pHM*Rluc* reporter expressing *Renilla* luciferase. Transfections also contained a firefly luciferase reporter vector lacking an RmRE to control for differences in DNA uptake or expression (6). Cell lysates then were assayed for *Renilla* and firefly luciferase activities relative to lysates derived from transfections lacking SP expression vectors (pcDNA3). The results indicated that the BAP-tagged GFP-SP construct was functional, but had slightly lower activity compared to GFP-SP (Fig. 1C).

To directly test the SP retrotranslocation assay, we co-transfected a negative control vector (pcDNA3) or expression vectors for GFP-SP lacking a BAP tag or GFP-SP-BAP together with the BirA expression plasmid. Although a low level of biotin is present in normal cells (37), it is usually insufficient to allow efficient biotinylation by BirA. Therefore, after 24 h, we added biotin to the media for 12 h, and cell lysates were prepared. Streptavidin was added to one portion of each extract, and treated and untreated extracts were analyzed by gel electrophoresis and Western blotting using GFP-specific antibody (Fig. 1D). As expected, lysates transfected with pcDNA3 alone lacked GFP-SP expression (lanes 1 and 2). Samples derived from transfection of the GFP-SP expression vector lacking a BAP tag showed a band consistent with GFP-SP, a product of signal peptidase cleavage, both in the presence or absence of streptavidin (compare lanes 3 and 4). As expected, lysates expressing GFP-SP-BAP revealed a band with a slightly higher mass than untagged GFP-SP. In the presence of streptavidin, we observed that a portion of GFP-SP-BAP was retarded within the gel, consistent with the tight binding of streptavidin to the biotinylated form of the protein (Fig. 1D, compare lanes 5 and 6). Only a portion of GFP-SP-BAP was biotinylated due to the addition of biotin after SP retrotranslocation to the cytoplasm and prior to its import into the nucleus for MMTV RNA binding (1). As expected, SP tagged at the N-terminus (see Fig. 1B), was biotinylated by the cytosolic BirA as detected by streptavidin binding, (Fig. 1D, compare lanes 7 and 8). These data strongly suggest that the C-terminal BAP tag of GFP-SP-BAP is biotinylated by BirA after retrotranslocation into the cytosol.

To verify that SP biotinylation occurred in transfected cells and not after lysis, we transfected HEK 293 cells with the GFP-SP-BAP expression plasmid together with the BirA expression vector or cells were transfected separately with each vector. After addition of biotin, lysates from cells individually transfected with the expression vector for GFP-SP-BAP or BirA were combined and compared to lysates from co-transfected cells. Streptavidin was added to an equal portion of each lysate prior to gel electrophoresis and Western blotting with antibodies specific for GFP and GAPDH (Fig. 1E). Lysates obtained from cells co-transfected with expression vectors for GFP-SP-BAP and BirA revealed a biotinylated band in the presence of streptavidin, whereas the combined lysate from separately transfected cells did not (Fig. 1E, compare lanes 2 and 4, respectively). These results indicated that biotinylation of GFP-SP-BAP occurs inside transfected cells, and not after cell lysis.

To confirm the SP membrane orientation and specificity of the retrotranslocation assay, we used a BirA expression construct engineered with an N-terminal signal peptide and a C-terminal KDEL sequence to allow expression and retention in the ER lumen (ER-BirA) (38). Cells were transfected with N-terminally or C-terminally BAP-tagged SP expression plasmids (GFP-BAP-SP or GFP-SP-BAP, respectively) and vectors expressing either BirA or ER-BirA. After addition of biotin, lysates were prepared and equal amounts were incubated with or without streptavidin prior to Western blotting. As anticipated for N-terminally BAP-tagged SP, biotinylated protein was detected in the presence of BirA and streptavidin (Fig. 1F, lane 2), but not when ER-BirA was expressed (lane 4). In contrast, biotinylated SP was detected when GFP-SP-BAP was expressed in the presence of ER-BirA (Fig. 1F, compare lanes 4 and 6). Interestingly, C-terminally BAP-tagged SP appeared to be more stable relative to the N-terminally BAP-tagged protein (Fig. 1F, compare lanes 5 and 6 and lanes 1-4). This result is consistent with the lower functional activity of GFP-BAP-SP in reporter assays (Fig. 1C). Together, our experiments support the type II transmembrane orientation of SP and verify the specificity of SP retrotranslocation assays.

### SP retrotranslocation requires the VCP/p97 ATPase

We previously have shown that SP function in a transient assay requires both the nuclear localization signal on the SP expression plasmid and the presence of the RmRE on the pHM*Rluc* reporter vector (1). Using this assay, we demonstrated that a dominant-negative form of p97/VCP inhibited SP activity, but not Rem cleavage by signal peptidase (2). To confirm that our retrotranslocation assay was dependent on p97, we used a chemical inhibitor (CB-5083) that binds selectively to the D2 domain of the ATPase (39). HEK 293T/17 cells were transfected with vectors expressing GFP-SP-BAP and BirA for 24 h. Media containing 1 μM of inhibitor was added for 8 h prior to addition of biotin. We then treated an equal amount of each lysate with streptavidin before analysis by denaturing gel electrophoresis and Western blotting with GFP-specific antibody (Fig. 2A). Use of GAPDH-specific antibody confirmed equal loading of the lanes. Gels then were scanned and retrotranslocation was quantified. The results revealed that inhibition of p97 ATPase activity interfered with SP retrotranslocation (Fig. 2A, compare lanes 2 and 4).

**FIG 2.**
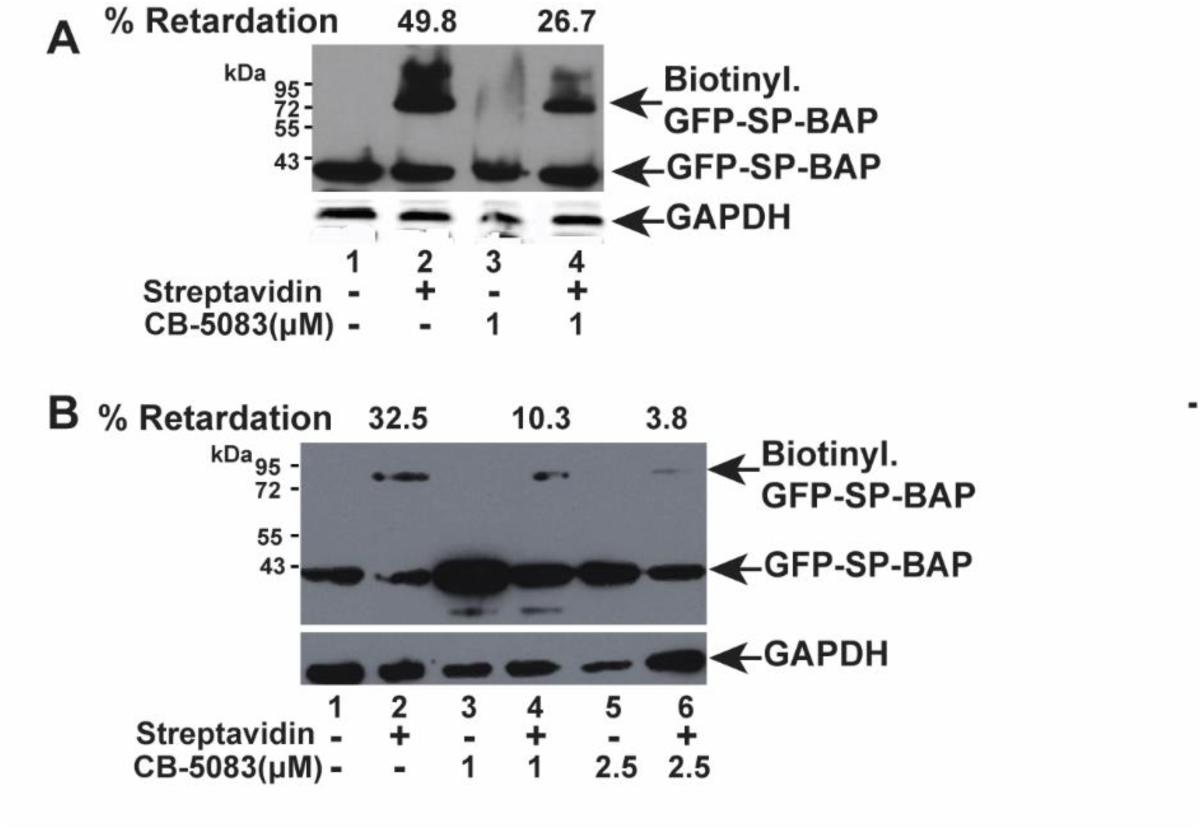
VCP/p97 is essential for SP retrotranslocation. (A) The VCP/p97 inhibitor CB-5083 reduces SP retrotranslocation in HEK 293T/17 cells. Cells were co-transfected with plasmids expressing GFP-SP-BAP and BirA. After 24 h, cells were treated with 1 μM CB-5083 for 8 h prior to addition of 100 μM biotin for 1 h. Cell lysates were incubated in the presence and absence of streptavidin prior to Western blotting and incubation with GFP or GAPDH-specific antibodies. The amount of biotinylated GFP-SP-BAP was quantitated relative to the total GFP-SP-BAP signal and given as % retardation. (B) CB-5083 blocks SP retrotranslocation in HC11 mouse mammary cells. The inhibitor was added at 24 h post-transfection for 9 h at the indicated concentrations. Biotin was added in the presence of CB-5083 for 1 h prior to removal of cells for lysate preparation. Streptavidin was added to one portion of each lysate and subjected to Western blotting. Blots were incubated with antibodies specific for GFP or GAPDH as indicated. The amount of biotinylated GFP-SP was quantitated relative to the total SP signal (% retardation).

Retrotranslocation assays with the VCP/p97 inhibitor also were performed in HC11 mouse mammary cells, a cell type that is a physiological target of MMTV infection (40). HC11 cells were transfected with an expression vector for GFP-SP-BAP together with the BirA expression plasmid and treated with either of two different amounts of CB-5083 for 9 h. Lysates were analyzed with and without streptavidin (Fig. 2B). As expected, quantitation of the results revealed that CB-5083 inhibited SP retrotranslocation in a dose-dependent manner (Fig. 2B, compare lane 2 to lanes 4 and 6). Since CB-5083 blocks VCP enzyme function, these results confirmed that p97 ATPase activity was required for SP retrotranslocation in both human and mouse cells.

### Biotinylated SP is sensitive to trypsin during retrotranslocation

To confirm that SP biotinylation is specific to proteins that have been extracted from the ER membrane, we used a trypsin-sensitivity assay with cell lysates containing intact microsomes. In this assay, we anticipate that SP extracted from the membrane and biotinylated in the cytosol will be susceptible to digestion with trypsin. In contrast, proteins that reside inside membranes are expected to resist trypsin digestion. HEK 293T/17 cells co-transfected with plasmids expressing C-terminally BAP tagged SP (GFP-SP-BAP) and BirA were harvested and subjected to freeze-thaw in a hypotonic buffer to isolate intact ER-derived microsomes. Lysates then were treated with increasing concentrations of trypsin. Streptavidin was added to one of the duplicate samples prior to gel electrophoresis and Western blotting. As a positive control for isolation of intact ER-derived vesicles, we used antibody against BiP, a cellular ER-resident protein (41). Samples treated with different concentrations of trypsin or with streptavidin showed little difference in the levels of BiP (Fig. 3A). These data confirmed that proteins within the ER lumen were protected from trypsin digestion under these conditions.

**FIG 3.**
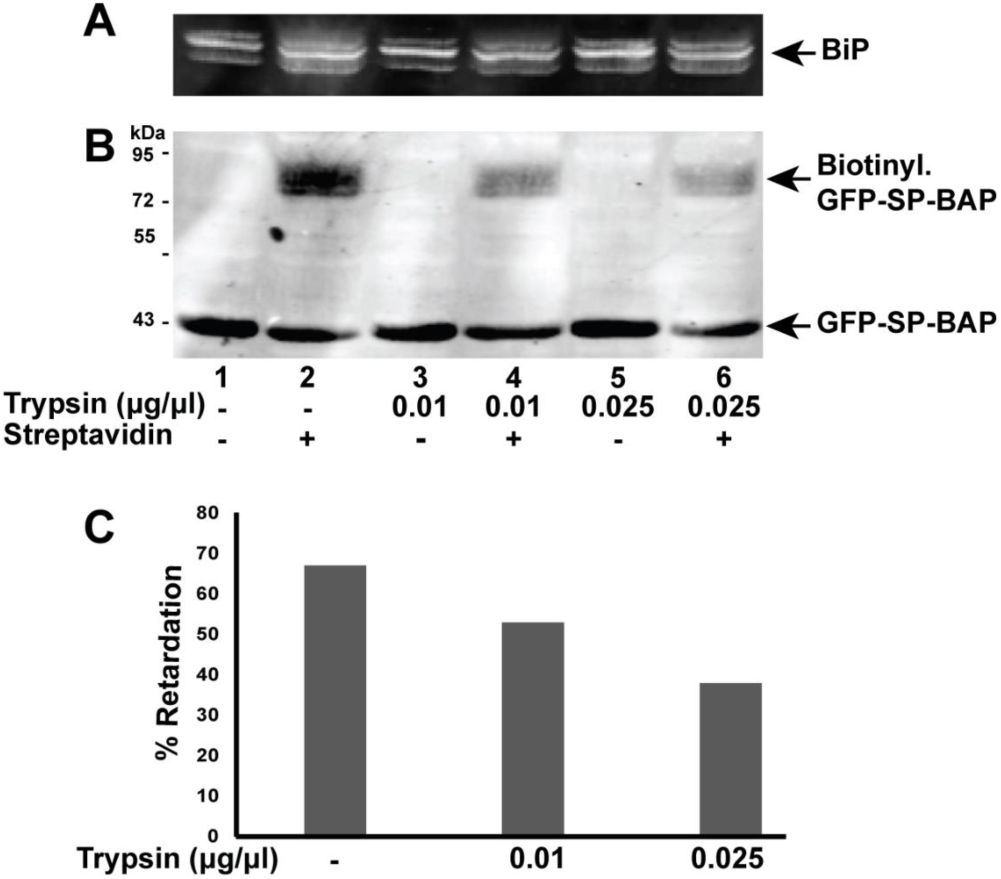
Trypsin sensitivity assay confirms that retrotranslocated SP is susceptible to enzymatic digestion. (A) BiP levels provide a measure of intact microsomes. Microsomes isolated from HEK 293T cells co-transfected with vectors for GFP-SP-BAP and BirA were treated with increasing trypsin concentrations. Western blots of microsome preparations were analyzed with BiP-specific antibody. (B) Biotinylated SP is trypsin sensitive. Microsomes were treated with the indicated concentrations of trypsin prior to addition of streptavidin and Western blotting with GFP-specific antibody. The majority of SP is localized in the nucleus (1) and is protected from trypsin digestion. (C) Quantitation of retrotranslocated GFP-SP-BAP in the presence of increasing trypsin concentrations. The level of biotinylated GFP-SP was quantitated relative to the total GFP signal and shown as percentage of retardation in the presence of streptavidin.

To determine whether biotinylated GFP-SP was susceptible to trypsin digestion, we incubated the blot with GFP-specific antibody. A significant portion of the signal in the absence of trypsin was shifted in the presence of streptavidin (Fig. 3B, compare lanes 1 and 2). However, the biotinylated fraction of GFP-SP-BAP showed a gradual decline with increasing trypsin concentration, indicating that the biotinylated GFP-SP is exposed to the cytosol. Quantitation of the trypsin sensitivity of the biotinylated protein confirmed this result (Fig. 3C). However, the non-biotinylated GFP-SP was trypsin resistant, suggesting that GFP-SP synthesized prior to the addition of biotin is protected from enzymatic digestion by the presence of a membrane. Since the steady state levels of SP are highest within nucleoli (1), these results suggest that non-biotinylated GFP-SP is protected from trypsin by the nuclear membrane.

### SP co-immunoprecipitation with VCP/p97 is ATP-dependent

Our previously published work indicates that VCP is required for efficient SP function (2, 21). We also have shown that VCP/p97 was identified in a covalent interaction screen using an SP-ubiquitin-activated interaction trap (SP-UBAIT) (21), and that a p97 inhibitor interfered with SP biotinylation in a retrotranslocation assay (see Fig. 2). Therefore, our expectation was that VCP interacts with SP during retrotranslocation and that p97-specific antibody would immunoprecipitate SP.

HEK 293 cells were co-transfected with expression vectors for GFP-SP and, in some cases, VCP/p97. After 48 h, cells were fixed in formaldehyde to preserve protein interactions, and a portion of the cell lysate was used for co-immunoprecipitation with p97-specific antibody. Immunoprecipitates and cell lysates were analyzed by Western blotting with p97 and GFP-specific antibodies. Little difference was observed with or without exogenous p97 expression (Fig. 4, right panel). GFP-SP was precipitated both in the presence and absence of exogenous p97 (Fig. 4, left panel). However, SP co-immunoprecipitation was increased when ATP and Mg^2+^ were added to the lysates (Fig. 4, compare lanes 2 and 3 to lanes 4 and 5). This result was not unexpected since the N-terminal domain of p97 is known to change its conformational state dependent on ATP binding (42). These experiments confirm that SP interacts with endogenous p97. In addition, our data suggest that SP binds to the ATP-bound form of VCP/p97.

**FIG 4.**
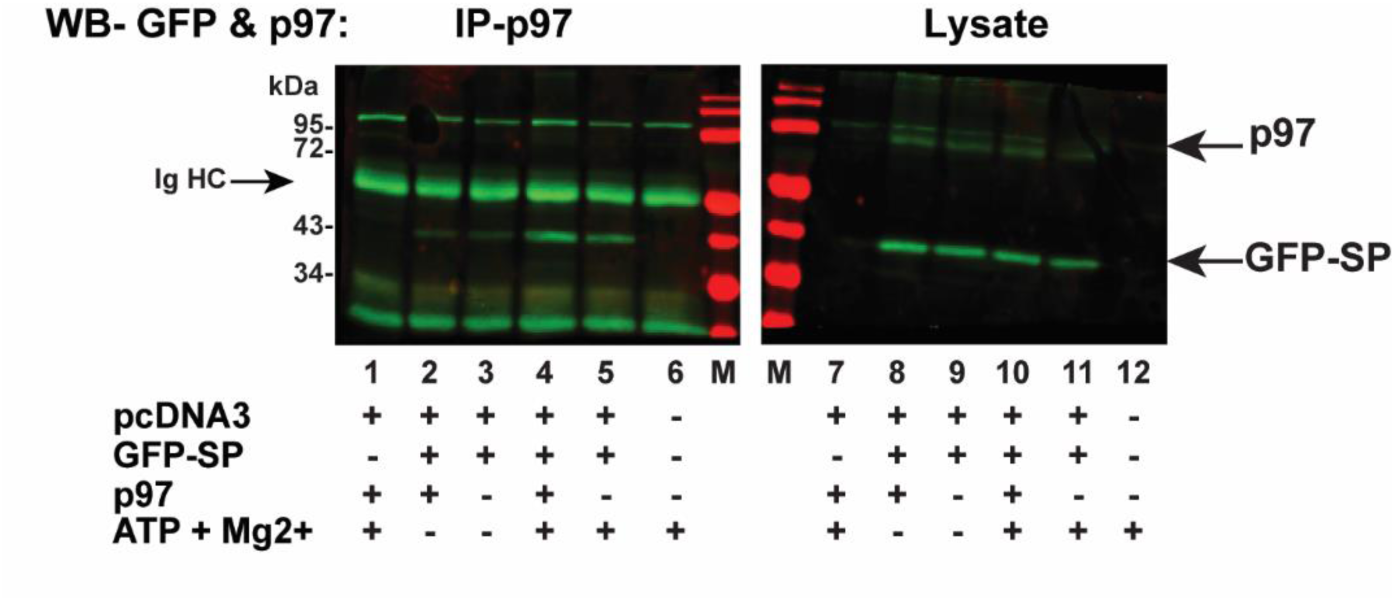
SP interaction with VCP/p97 is ATP-dependent. Cells (HEK 293) were transfected with pcDNA3 and an expression plasmid for GFP-SP in the presence or absence of an exogenous VCP/p97 expression vector. Co-mmunoprecipitation of cell lysates with p97-specific antibodies was performed with or without exogenous ATP and its cofactor Mg^2+^ as indicated. Western blotting with GFP-specific antibody is shown for co-immunoprecipitates (lanes 1-6) or cell lysates (lanes 7-12). M = molecular mass markers. The positions of immunoglobulin heavy chain (Ig HC), p97, and GFP-SP are indicated with arrows.

### SP amino acids 57-58 are required for retrotranslocation and efficient interaction with p97

To identify SP regions essential for the retrotranslocation process, we prepared a series of N-terminal deletion mutants (Fig. 5A), including the nuclear/nucleolar sequence (NLS/NoLS) (1, 7). Our previous experiments have shown that the NLS/NoLS is required for SP function in reporter assays (1). Plasmids expressing GFP-SP-BAP or SP deletion mutants and BirA were co-transfected into HEK 293 cells. Transfected cells were treated with biotin prior to extract preparation. Extracts were subjected to Western blotting in the presence or absence of streptavidin. Deletion of the N-terminal 50 amino acids of SP within the GFP-SP-BAP construct retained SP retrotranslocation activity (Fig. 5B). Deletion of the N-terminal 60 amino acids of SP produced an unstable protein despite the presence of the GFP tag (Fig. 5B, lanes 13 and 14). Biotinylation assays using these mutants suggested that the SP region C-terminal to amino acid 50 was important for retrotranslocation (Fig. 5B, even lanes 4-12 compared to 14). We also assayed the functional activity of the deletion mutants. Cells were transfected in triplicate with expression vectors for wild-type and SP mutants in addition to the pHM*Rluc* reporter plasmid. In contrast to retrotranslocation assays, N-terminal deletions of 30 amino acids or more eliminated detectable SP function (Fig. 5C), consistent with removal of a portion of the NLS/ARM (Fig. 5A).

**FIG 5.**
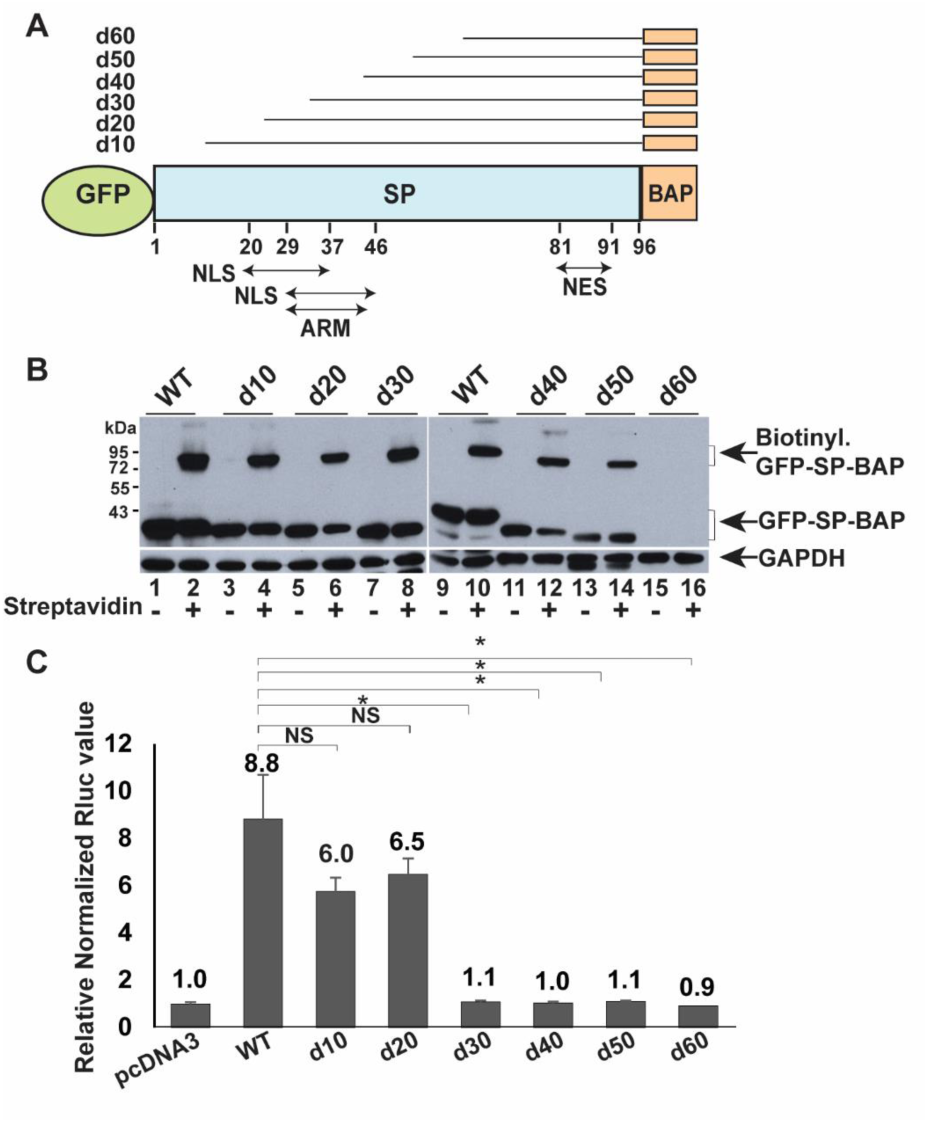
The N-terminal 50 amino acids of SP, including the NLS, are not required for retrotranslocation. (A) Diagram of N-terminal SP deletions. Each of the mutants is tagged at the N-terminus and C-terminus with GFP and BAP, respectively. Sequential increments of 10 amino acids were deleted from the N-terminus of SP (shown as d10-d60). The positions of the NLSs, ARM, and NES are indicated. Two amino acids of the SP C-terminus were deleted to prevent cleavage of the BAP tag by signal peptidase. (B) Retrotranslocation assays with SP deletion mutants. Cells (HEK 293) were co-transfected with expression vectors for BirA and GFP-SP-BAP or deletion mutants. After addition of biotin, lysates were used for streptavidin addition and Western blotting with GFP and GAPDH-specific antibodies. The position of biotinylated and non-biotinylated GFP-SP proteins are shown with arrows. (C) Reporter assays for GFP-SP-BAP deletion mutants. Cells were co-transfected in triplicate with the reporter vector pHM*Rluc* and expression vectors for wild-type GFP-SP-BAP or deletion mutants. Results were reported as described in Fig. 1C. NS = not significant. * = p<0.05.

To avoid potentially destabilizing effects of large deletions, we performed alanine-scanning mutagenesis to delineate further SP sequences needed for ER membrane extraction. We prepared two-to-three amino-acid substitutions in the region between 50 to 61 amino acids, which is localized between the NLS/NoLS/ARM and the NES (Fig. 6A). Co-transfections of plasmids expressing wild-type SP or mutants and BirA were used for biotinylation in HEK 293 cells. Extracts incubated with streptavidin showed that mutants M50-52, M53-55, and M59-61 had retrotranslocation efficiencies similar to wild-type GFP-SP-BAP (Fig. 6B). Since wild-type SP has an alanine at position 56, only two amino acids were mutated for M57-58 (WQ to AA). Alteration of these two amino acids decreased retrotranslocation activity, which varied depending on the efficiency of transfection and the amount of expression plasmid used (Fig. 6C). In multiple experiments (n = 9) subjected to quantitation relative to the total GFP signal detected, M57-58 showed significantly reduced retrotranslocation compared to wild-type SP (Fig. 6D).

**FIG 6.**
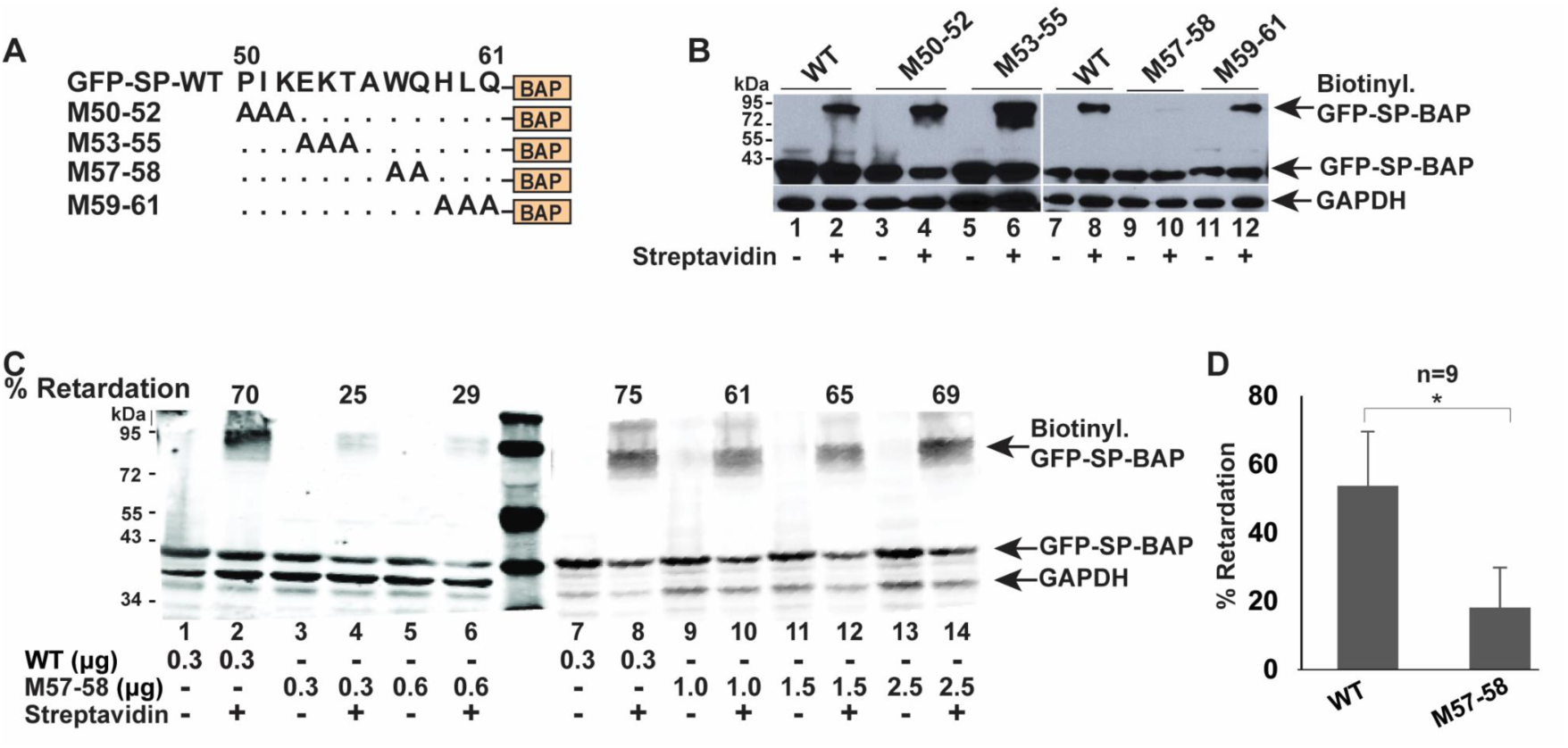
Amino acids 57-58 of SP are critical for retrotranslocation. (A) Diagram and nomenclature for sequential alanine substitution mutations in GFP-SP-BAP starting from SP amino acid 50. (B) Retrotranslocation assays with alanine substitution mutants. HEK 293T cells were co-transfected with expression vectors for BirA and wild-type or mutant GFP-SP-BAP. Biotin was added for 12 h prior to extract preparation and addition of streptavidin in alternate lanes. Western blots were incubated with GFP and GAPDH-specific antibodies. Arrows indicate the positions of biotinylated and non-biotinylated GFP-SP. (C) Retrotranslocation assay with increasing levels of mutant M57-58 compared to wild-type SP. Transfections were performed as in panel B. The amounts of the wild-type and mutant expression vector used for transfection are provided in micrograms. The % retardation is calculated by dividing the signal of biotinylated GFP-SP compared to the total GFP signal. (D) Quantitation of multiple retrotranslocation assays comparing wild-type GFP-SP-BAP and M57-58. Transfections of HEK 293T cells were performed with 300 and 600 ng of wild-type and mutant SP expression vectors, respectively, to ensure equal SP levels together with the BirA expression plasmid. Biotin was added from 2 to 12 h prior to lysate preparation in different experiments. The means and standard deviations of nine quantitated assays showed that M57-58 has a significantly decreased retrotranslocation efficiency compared to the wild-type. * = p<0.05.

The M57-58 defect in retrotranslocation predicted that this mutant would be defective in the SP reporter assay. Therefore, we tested substitution mutants for SP activity. Cells were co-transfected in triplicate with expression vectors for GFP-SP-BAP (wild-type) or mutants together with the pHM*Rluc* plasmid. As anticipated, no activity was detected for M57-58 using the same amount of the expression vector that gave a 6-fold increase in luciferase activity for the wild-type plasmid (Fig. 7A). No statistical difference was observed between the wild-type SP and the mutants, M50-52 and M53-55. Surprisingly, the M59-61 mutant, which showed no defect in the retrotranslocation assay, had greatly reduced SP reporter activity compared to the wild-type expression vector. We repeated the activity assays for both M57-58 and M59-61 with higher concentrations of the expression plasmids (Fig. 7B). The M57-58 mutant showed minimal activity even at 5-fold higher amounts of DNA in the reporter assay, but remained significantly lower than wild-type SP. The M59-61 mutant had detectable activity using DNA amounts that were significantly more compared to the wild-type expression vector (Fig. 7B). These data suggest that M59-61 has a defect in a function subsequent to SP retrotranslocation to the cytosol.

**FIG 7.**
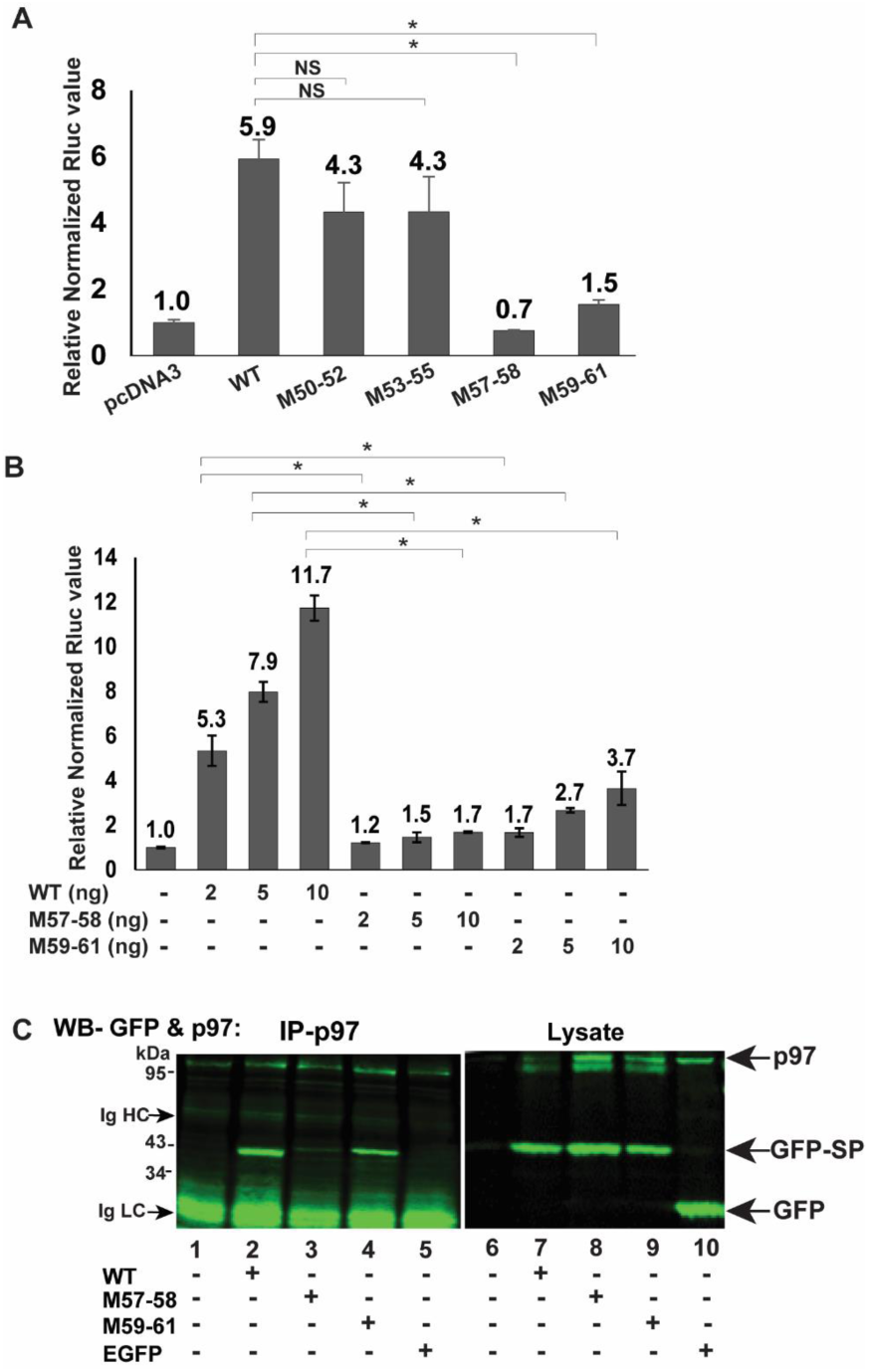
Amino acids needed for p97 interaction are a subset of sequences required for optimal SP function. (A) Mutants M57-58 and M59-61 are defective for SP function. HEK 293T cells were co-transfected in triplicate with expression vectors for GFP-SP-BAP (WT) alanine scanning mutants (2.5 ng each), the SP reporter vector pHM*Rluc* (250 ng), and pGL3-control (250 ng). Luciferase values were normalized for pGL3-control expression in each transfection and expressed relative to transfections with pcDNA3 lacking SP expression (assigned a relative value of 1). Two-tailed Student’s t tests were performed (* p<0.05). NS = not significant. (B) Titration of M57-58 and M59-61 functional activity compared to WT GFP-SP-BAP. Expression vectors (DNA levels given in nanograms) were co-transfected with pHM*Rluc* reporter and pGL3-control as described in panel A. (C) Co-immunoprecipitation of GFP-SP and mutants with VCP/p97. HEK 293T cells were transfected with expression vectors for GFP, GFP-SP (WT), or mutants. Extracts were immunoprecipitated with antibody specific for p97. Immunoprecipitates (left panel) or cell lysates (right panel) were analyzed by SDS-polyacrylamide gel electrophoresis before Western blotting with antibodies specific for p97 and GFP. The positions of p97, GFP-SP, GFP, and immunoglobulin heavy and light chains are indicated by arrows.

Since SP extraction from the ER membrane is likely to be dependent on p97 based on the involvement of the ATPase in ERAD (43), we performed co-immunoprecipitation experiments with lysates obtained from cells transfected with plasmids expressing wild-type GFP-SP or the GFP-tagged versions of the mutants M57-58 and M59-61 (Fig. 7C). Precipitates and cell lysates were subjected to Western blotting with antibodies to p97 and GFP. Antibody specific for VCP/p97 precipitated both the wild-type SP as well as M59-61. However, the mutant M57-58 interacted very weakly with p97 in cellular lysates containing ATP and Mg^2+^ under the same conditions (compare lanes 2 and 3). Co-immunoprecipitations with lysates from cells transfected with EGFP expression vector served as a negative control (Fig. 7C, lane 5). Analysis of cell lysates revealed similar expression of the wild-type and mutant SP proteins (Fig. 7C, lanes 7-9). Furthermore, GFP alone did not immunoprecipitate with p97 (lane 5). Our experiments suggest that SP amino acids 57 and/or 58 (WQ) are essential for interaction directly or indirectly with the cellular ATPase p97.

### SP mutants 57-58 and 59-61 show aberrant nuclear localization

SP mutants M57-58 and M59-61 had decreased activity in our reporter assay and amino acids 57-58 appear to be critical for SP retrotranslocation. Therefore, we anticipated increased co-localization of M57-58 with a fluorescent ER marker, ER-mCherry, compared to wild-type SP. ER-mCherry has an N-terminal signal peptide linked to monomeric Cherry and a C-terminal KDEL to facilitate retention in the ER lumen (44). HC11 mouse mammary cells were co-transfected with plasmids expressing ER-mCherry and either wild-type GFP-SP or SP mutants prior to fixation, nuclear staining with 4′,6-diamidino-2-phenylindole (DAPI), and confocal microscopy (Fig. 8A). Our published observations have shown that wild-type SP is localized primarily in nucleoli similar to other HIV-1 Rev-like proteins (19, 20), in agreement with the results observed here (Fig. 8A; see top right, WT Merge). In contrast, M57-58 appeared in perinuclear regions that overlapped with the ER-mCherry signal as well as the nucleus. M59-61 was co-localized with the mCherry signal, but also outside the ER in both nuclear and extranuclear locations. Compared to the wild-type GFP-SP, neither mutant appears to localize to the nucleolus (Fig. 8A, middle and lower panels). Quantitation of the fluorescent signals in multiple cells from two independent experiments revealed some wild-type SP co-localized with the ER marker (Fig. 8B), which is consistent with SP synthesis in the ER prior to retrotranslocation and nuclear trafficking. However, the mutant M57-58 showed low, but significantly higher, co-localization with the ER marker relative to wild-type SP (0.03 and 0.00, respectively) (Fig. 8B). The mutant M59-61, which had normal activity in the retrotranslocation assay, showed a similar localization pattern as M57-58 (Fig. 8A and 8B), indicating that M59-61 also had defective nucleolar trafficking. These results explain the low activity of M57-58 and M59-61 in reporter assays, which depend on SP binding to the RmRE, presumably in nucleoli.

**FIG 8.**
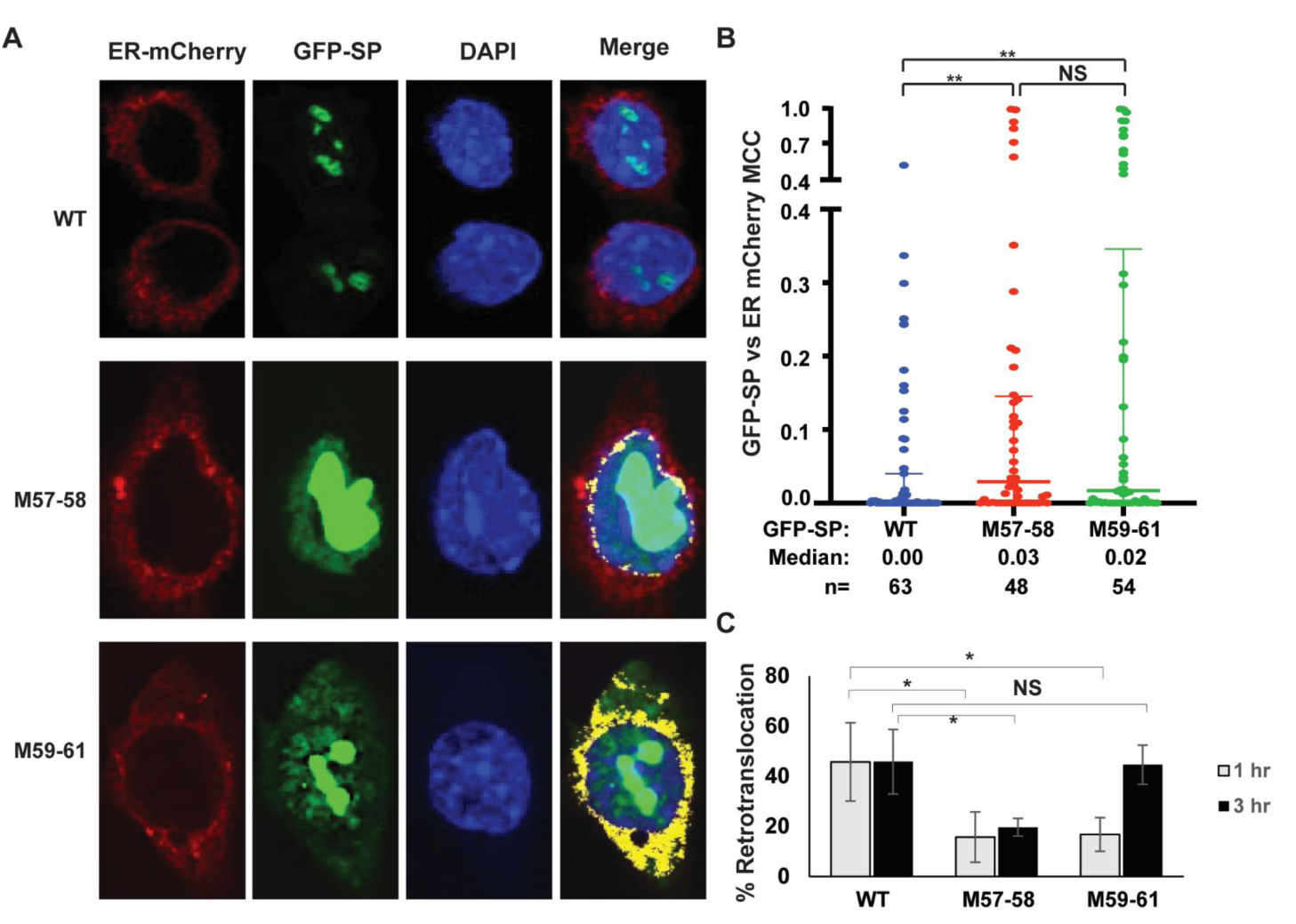
SP mutants 57-58 and 59-61 show increased ER co-localization and reduced retrotranslocation relative to wild-type SP. (A) Colocalization of wild-type or mutant SP with an ER marker in individual cells. HC11 mammary cells were co-transfected with expression vectors for GFP-SP (WT) or mutants (green) together with ER-mCherry (red). Cells were fixed, and the nuclei were stained with DAPI (blue). A merged image is provided on the right. Yellow areas in the merged images indicate red and green overlapping pixels determined by the colocalization analysis using ZEN software. Cells expressing GFP-SP (WT) lack colocalization between the red and green signals, so no yellow areas were observed. (B) Quantitation of the GFP-SP signal colocalization with ER-mCherry. Each dot represents the Mander’s colocalization coefficient (MCC) of GFP-SP with the ER signal for an individual cell as suggested by Manders under conditions in which signal intensities are unequal (73). The median signal was calculated based on the numbers of cells analyzed (n). Mann-Whitney t tests were used to compare wild-type and mutant MCCs. ** = p<0.01; NS = non-significant. (C) Retrotranslocation assays with pulses of biotin indicate defects in the ER membrane extraction of both M57-58 and M59-61. HEK 293T cells were co-transfected with 300 ng of expression vectors for wild-type GFP-SP-BAP or M59-61 or 600 ng of the M57-58 expression vector together with a BirA expression plasmid. Cells were pulsed for either 1 h or 3 h prior to lysate preparation and Western blotting. Retrotranslocation was quantified as described in the Fig. 2 legend. The results of triplicate assays were expressed as the means and standard deviations for the 1 h and 3 h biotin pulses. Student’s t tests were performed between the wild-type and mutants. The asterisk shows a p-value of <0.05, whereas NS indicates a non-significant value.

Our previous retrotranslocation assays were performed with lysates obtained from transfected cells treated with biotin for 4 to 12 h to maximize detection of SP extraction from the ER membrane. To determine the timing of the retrotrotanslocation process, we performed shorter pulses of biotin after co-transfection of HEK 293T cells with plasmids expressing the wild-type SP (GFP-SP-BAP) or the mutants with a BirA expression vector. At 24 h post-transfection, cells were treated with 100 μM biotin for either 1 h or 3 h. Lysates were incubated with streptavidin and used for Western blotting with GFP-specific antibody prior to quantitation (Fig. 8C). Retrotranslocated SP constituted approximately 40 to 50% of the total wild-type SP signal and was similar for lysates obtained at 1 h and 3 h after biotin addition. As expected, the M57-58 mutant showed <20% retrotranslocation during the 1 h-biotin treatment, and the longer incubations did not significantly change this result. In contrast, the M59-61 mutant showed an initial defect in retrotranslocation that was detectable after 1 h of biotin treatment, whereas retrotranslocation of this mutant recovered to wild-type SP levels after 3 h. Together with previous results, our data suggest that both M57-58 and M59-61 have defects in retrotranslocation to different extents (Fig. 7C). Nevertheless, M57-58 has a weaker interaction with VCP/p97, which explains its greater defect in ER membrane extraction.

### Conformational modeling predicts defective multimerization of SP mutants

SP is a Rev-like protein that likely shares a similar secondary structural conformation. To test this idea, we performed molecular modeling of SP using AlphaFold2, a highly sophisticated machine learning-based modeling tool (45, 46). SP is predicted to contain a long disordered N-terminus and three α-helices designated as α1, α2 and a transmembrane helix (as detected by MEMSAT-SVM (47) (Fig. 9A). Since SP is a type II transmembrane protein, the disordered N-terminus faces the cytosol, and the very short, disordered C-terminus localizes within the ER. The α1 helix contains the NLS/ARM and is connected to the α2 helix by a loop. This structure is reminiscent of HIV-1 Rev, which is known to form a helix-loop-helix (helical hairpin) conformation. The α-helices of HIV-1 Rev are known to form a coiled-coil structure that acts as a nucleation site for oligomerization (48).

**FIG 9.**
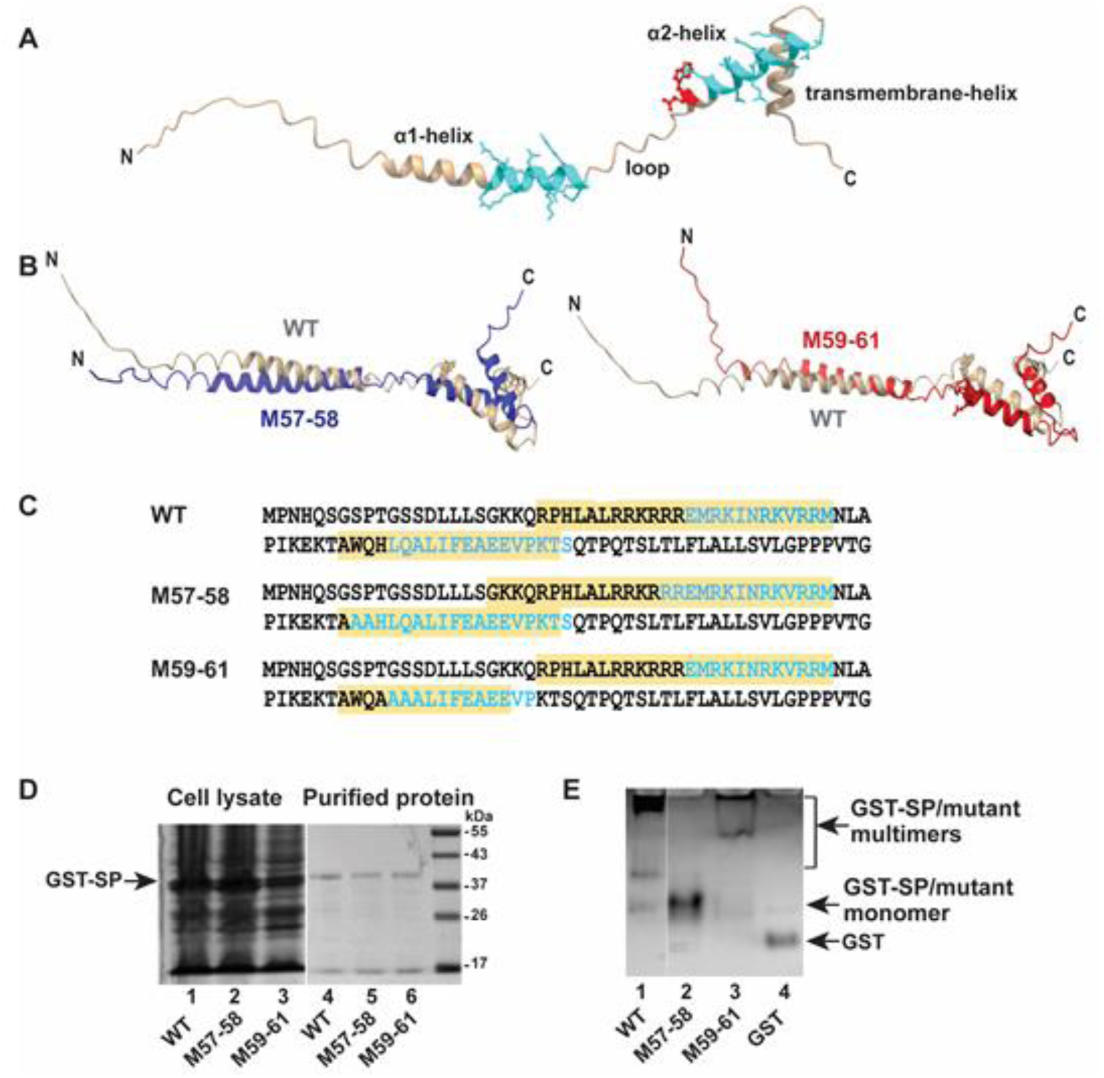
Structural models of SP mutants predict conformational changes related to wild-type SP and reduced multimerization in native gels. A) Molecular modeling of SP using AlphaFold2. The two α-helices (α1 and α2) and the transmembrane helix are indicated; α1 and α2 are separated by a loop. The coiled-coil motifs in SP as predicted by the CCHMM_PROF coiled-coil server are shown in aqua. The p97-interacting residues W57 and Q58 are depicted in red. The model was analyzed and represented using ChimeraX. (B) Predicted α-helices in wild-type relative to mutant SP. Wild-type (WT), M57-58 and M59-61 have been represented by gray, blue, and red, respectively. M57-58 has an extended α1-helix (left image), whereas M59-61 has a shortened α2-helix (right image). (C) Amino acid sequence of WT and mutants comparing the α-helical regions and coiled-coil motifs. The sequence of α1 and α2 helices are highlighted in yellow, whereas those forming the coiled-coil motifs are depicted in aqua. M57-58 is predicted to have an increased length of both coiled-coils. The second coiled-coil formed by the α2-helix is predicted to be shorter for M59-61. (D) Integrity of purified GST-tagged versions of wild-type and SP mutants. The purity and integrity of purified proteins were confirmed by analysis on a denaturing SDS-containing polyacrylamide gel. Cell lysates have been shown on the left and compared to purified proteins on the right. The position of GST-SP is shown with an arrow. (E) Gradient gel electrophoresis of purified WT GST-SP or M57-58 and M59-61 mutants under native conditions. Purified proteins (5 μg) were analyzed on a 4-20% Tris-buffered gradient gel lacking SDS. The relative positions of multimers and monomers as well as the unfused GST protein are shown with arrows.

The coiled-coil motifs in SP were modeled using the CCHMM_PROF coiled-coil server (49). Like Rev, SP was predicted to have two coiled-coil motifs localized in the α1 and α2 helical regions. Since M57-58 and M59-61 differ from WT SP and each other, similar modeling predicted changes within the α-helices. The α1-helix of M57-58 was extended with no major conformational change in the α2-helix (Fig. 9B, left). However, mutation in the 59-61 region destabilized the α2-helix (Fig. 9B, right). Neither mutation affected the conformation of the transmembrane helix (Fig. 9B and 9C). Conformational changes in either α-helix could affect SP stability and function. Likewise, both predicted coiled-coils were extended in M57-58, whereas the mutation in M59-61 shortened the coiled-coil of the α2-helix (Fig. 9C). If SP forms the helical hairpin for RNA binding, changes in the α-helices and/or length of coiled-coil region might create stearic hindrance to interfere with the formation of oligomers/multimers.

To determine whether amino acid changes in M57-58 and M59-61 affect the ability of SP to form multimers, we expressed the wild-type and mutant SP proteins fused to glutathione-S-transferase (GST) in *E. coli*. Cell lysates were used for purification and analysis on SDS-containing gels prior to staining. The results revealed similar yield and purity of the proteins (Fig. 9D). To assess the ability of purified SP proteins as well as GST to oligomerize, we resolved the purified wild-type or mutant proteins using polyacrylamide gradient gel electrophoresis in the absence of denaturing agents. After Coomassie blue staining, the wild-type SP protein showed a band expected of the GST-SP monomer (Fig. 9E, lane 1), although most of the protein was located near the gel origin, indicative of higher oligomeric conformation. As expected, the purified GST control did not oligomerize (lane 4). In contrast, the majority of M57-58 appeared to be monomeric, whereas M59-61 migrated between the monomers and oligomeric wild-type SP. These experiments suggested that both M57-58 and M59-61 are defective in multimerization.

## DISCUSSION

Unlike other signal peptides, MMTV-encoded SP is unusually long (98 amino acids) and has an arginine-rich motif typical of RNA-binding proteins, an NLS/NoLS, and a leucine-rich NES typical of proteins that use the Crm1 export pathway (50–52). We previously showed that the NLS is required for SP localization to the nucleolus (1). SP must be extracted from the ER membrane by retrotranslocation prior to its entry into the nucleus for binding to the RmRE on MMTV RNA (6, 53). Retrotranslocation is a critical component of ERAD, which provides cellular quality control of protein function (24, 54, 55).

To directly measure SP retrotranslocation to the cytosol, we developed a quantitative assay. This assay is based on the biotinylation of a lysine residue in a C-terminally BAP-tagged SP protein during expression of the *E*.*coli* ligase BirA in the cytosol of mammalian cells. Similar assays have been used to measure the retrotranslocation of both membrane-bound and secretory proteins to the cytosol for ERAD (33, 34, 56). Our previous experiments indicate that SP is a type II transmembrane protein with the C-terminus in the ER lumen based on glycosylation of the Rem C-terminus (2). C-terminal BAP tagging of SP does not greatly compromise its function based on reporter assays (Fig. 1C). Biotinylation of the C-terminal BAP tag by the cytosolic BirA provides direct evidence that MMTV-SP is retrotranslocated from the ER to the cytosol. Moreover, this assay represents a simple, reliable, and quantitative measure of SP extraction from the ER membrane (Fig. 1D). The assay was validated by comparisons to functional reporter assays (Fig. 1C), the requirement for VCP/p97 ATPase activity (Fig. 2), the use of biotin ligases that were specific for either the cytosol or the ER lumen (Fig. 1F), and trypsin sensitivity of the biotinylated products not protected by the ER membrane (Fig. 3).

Multiple experiments support the idea that VCP/p97 is essential for efficient SP retrotranslocation. First, using our retrotranslocation assay, we confirmed that inhibition of p97 activity reduced SP extraction from membranes in two different cell lines (Fig. 2). Second, we showed that p97 interacted with MMTV-encoded SP by co-immunoprecipitation, and interactions were dependent on the presence of ATP and Mg2+ (Fig. 4). Our published results using a UBAIT screen support an SP-p97 interaction (21). Third, we identified a two amino-acid sequence (WQ) in mutant M57-58 that greatly decreased association with p97 (Fig. 6). Fourth, the same mutation that inhibited SP interaction with p97 showed a defect in retrotranslocation as well as SP function in our reporter assay (Fig. 7). Together with our previously published results that SP function is inhibited by expression of a dominant-negative p97 (2, 21), we conclude that VCP/p97 provides the energy required for extraction of SP from the ER membrane.

Although p97 is required for SP retrotranslocation, we cannot rule out the need for an adapter protein to mediate their association. Our previous high-throughput screen failed to identify any known p97 adapters (21), such as UFD1/NPL4 or UBX2, which often mediate interactions between ubiquitylated ERAD substrates and VCP/p97 (30, 57). Therefore, we speculated that SP interacts with p97 directly (21). However, our attempts to use binding assays with purified GST-tagged SP and His-tagged p97 purified from *E. coli* have not been successful. In support of our hypothesis, published data suggest that p97 associates with both ubiquitylated and non-ubiquitylated polypeptides (25, 58). In addition, the yeast homologue of p97, Cdc48 (30, 57, 59), has been shown to recognize monoubiquitylated substrates that are not degraded (60, 61). MMTV-encoded SP may mimic the p97 adapters, NPL4 and/or UFD1 (62, 63), to compete directly for p97 binding in the absence of ubiquitylation or an adapter may yet be identified.

The mutant M59-61, which showed a limited defect in retrotranslocation (Fig. 8C), had reduced activity in an assay for SP function (Figs. 7A and 7B). This mutant also showed normal interactions with p97 under the co-immunoprecipitation conditions used (Fig. 7C). Both M57-58 and M59-61 likely have defects in retrotranslocation as determined from limiting the biotinylation period to 1 h (Fig. 8C), but M57-58 is missing critical interactions with VCP/p97 that more dramatically hamper ER extraction (Fig. 7C). Confocal microscopy of both M57-58 and M59-61 showed slightly, but significantly, increased ER localization compared to wild-type SP and abnormal distribution at steady state (Fig. 8). Molecular modeling predicted that wild-type SP has three α-helices (Fig. 9A). Two of those helices likely form a coiled-coil structure. Furthermore, independent modeling of the functionally inactive mutants M57-58 and M59-61 revealed that each mutant changed the length of the α-helical and coiled-coil regions, predicting altered interactions (Figs. 9B and C). Direct testing of purified mutant proteins on non-denaturing gels confirmed this prediction, although M57-58 appeared to form less multimers than did M59-61 compared to wild-type SP (Fig. 9E). This result may reflect additional defects of M57-58 in membrane extraction that limit multimerization. Therefore, mutants in the SP region from 57-61 showed a slight increase in ER association and abnormal nuclear localization due to defects in interaction with p97 and adapters, and/or multimer formation.

In summary, MMTV-encoded SP interacts with a critical cellular factor, VCP/p97. Point mutations in VCP induce multi-system proteinopathy type 1 (MSP1), which is known as inclusion body myopathy associated with Paget’s disease and frontotemporal dementia/amyotrophic lateral sclerosis (IBMPFD/ALS) (27, 28, 64, 65). Disease mutations increase p97 ATPase activity and change interactions with p97 adapter proteins (62). Since MMTV SP is highly related to the human endogenous retrovirus type K (HERV-K) Rec protein (66–68), it is possible that Rec interacts with VCP/p97 to affect ERAD and normal protein quality control. Our results highlight the unique characteristics of long signal peptides specified by host and viral genes (13, 69). Many long signal peptides, including MMTV SP, are known to be signal recognition particle-dependent and are co-translationally transferred across the ER membrane (69). Nevertheless, these unique signal peptides are more than zip codes, suggesting that additional characterization will provide translational benefits for development of biotherapeutics (13).

## MATERIALS AND METHODS

### Constructs

The coding sequence of the BAP tag (GLNDIFEAQKIEWHE) (70) was inserted after amino acid 96 of N-terminally GFP-tagged SP followed by a stop codon using site-directed mutagenesis with the CloneAmp HiFi PCR Premix (Takara Bio). The BirA and ER-BirA constructs were obtained from Addgene (38, 71). The N-terminal deletion mutants of GFP-SP-BAP (d10-d60) were generated by PCR, and the products were cloned between the *Xho*I and *Bam*HI sites of GFP-SP. Alanine scanning mutagenesis was used to prepare specific GFP-SP-BAP mutants between SP amino acids 50 to 61. In this approach, two to three amino acids in GFP-SP-BAP were replaced by alanine for each construct using the CloneAmp HiFi PCR Premix (Takara Bio) according to the manufacturer.

### Cell lines and transfections

HEK 293, 293T, or 293T/clone 17 (293T/17) cells (purchased from ATCC) were cultured at 37°C and 7.5% CO_2_in Dulbecco’s modified Eagle’s medium (DMEM; Life Technologies, Inc.) supplemented with 10% fetal bovine serum (FBS), 100 U/ml penicillin, 100 U/ml streptomycin, 50 μg/ml gentamicin, and 2 mM L-glutamine. Cells were seeded in 6-well plates at 1 × 10^6^ cells/well for 24 h prior to transfection. Polyethylenimine (PEI) transfection reagent (72) was added with the indicated amounts of DNA and adjusted to a total of 3 μg of DNA with the plasmid vector pcDNA3. Most biotinylation experiments used 300 ng each of GFP-SP-BAP and BirA constructs. After 24 h, the media was replaced with 2 ml of serum-free media supplemented with 0.1 mM biotin and further incubated for 1 to 12 h as noted. In transfections using luciferase reporter vectors, 250 ng each of the pHM*Rluc* (1) and pGL3-control plasmids (Promega) were added in addition to the indicated amounts of SP expression constructs.

### Cell lysate preparation, biotinylation assay and Western blotting

Transfected HEK 293, 293T, or clone T17 cells were washed with phosphate-buffered saline (137 mM NaCl, 2.7 mM KCl, 10 mM NaH_2_PO4, and 1.8 mM KH_2_PO4) to remove free biotin. Cell pellets were lysed in SDS-sample buffer (25 mM Tris-HCl, pH 6.8, 1% SDS, 10% glycerol, and 5% 2-mercaptoethanol). For gel retardation assays, samples were boiled for 10 min, cooled to room temperature, and incubated with 1 μg of streptavidin (Sigma) for 30 min before separation on SDS-containing polyacrylamide gels. Proteins were then transferred to nitrocellulose membranes. Membranes were pre-incubated with Tris-buffered saline [20 mM Tris-HCl, pH 7.4, 137 mM NaCl (TBS)] plus 0.1% Tween 20 (TBS-T) buffer containing 5% nonfat dry milk (TBS-TM) for 1 h. Membranes were incubated with primary antibody in TBS-TM for 2 h at room temperature or overnight at 4°C, depending on the antibody. Following three washes, the membranes were treated with horseradish peroxidase-conjugated secondary antibody in TBS-TM for 1 h, and then washed again thrice with TBS-T. Secondary antibody binding was detected using the ECL Prime Western Blotting Detection Reagent (Amersham) as described by the manufacturer. For LI-COR detection, membranes were incubated with Odyssey Blocking Buffer (OBB) and TBS (1:2 ratio by volume) followed by incubation with primary antibody diluted in OBB–TBS-T (1:2 ratio by volume). The membranes were then washed thrice with TBS-T and incubated with fluorescently labeled IRDye secondary antibodies in OBB–TBS-T (1:2 ratio by volume) for 1 h. Following the incubation, membranes were washed again thrice in TBS-T and once in TBS. Signals were then detected using the Odyssey Imaging System according to the manufacturer’s instructions.

The sources of the antibodies were: GFP (Clontech); GAPDH, BiP, goat anti-mouse IgG, goat anti-rabbit IgG (all from Cell Signaling); VCP (Novus Biologicals or Invitrogen); IRDye 800CW goat anti-mouse IgG and IRDye 680RD goat anti-rabbit IgG (LI-COR). Band intensities were quantitated either using ImageJ or Image Studio software.

### Trypsin sensitivity assays

To maintain intact microsomes for trypsin sensitivity assays, cells were resuspended in a buffer containing 50 mM Tris-HCl, pH 8.0, 250 mM sucrose, and 10 mM N-ethyl-maleimide (to block BirA post-lysis activity). Cell suspensions were subjected to one freeze-thaw cycle and centrifugation at 5,000 Xg for 5 min at 4ºC. Supernatants (microsome-containing lysates) containing ∼100 μg of total protein were incubated with 0.01 μg or 0.025 μg trypsin in reaction volumes of 100 μl for 1 h at 37ºC. Microsome preparations then were used for retardation assays.

### Co-immunoprecipitation experiments

For co-immunoprecipitation studies, different GFP-tagged SP constructs (either wild-type or mutants) (1 μg) were used for transfection of HEK 293T/17 cells. After 48 h, cells were harvested and fixed in 4% formaldehyde solution. Cells were lysed in NP-40 lysis buffer (0.1 M Tris-HCl, pH 7.5, 0.1 M NaCl, 1% NP-40, and 1 mM dithiothreitol [DTT]) containing 1 mM ATP, 5 mM MgCl_2_, and complete protease inhibitor (Thermo Fisher Scientific) on ice for 1 h with intermittent vortexing. The lysates were then cleared by centrifugation at 13,000 Xg at 4ºC, and the protein yield was determined. Lysates containing ∼200 to 500 μg of total protein were incubated with rabbit polyclonal p97-specific antibody (Novus Biologicals) overnight at 4ºC. Antibody-containing mixtures were incubated with magnetic Protein G beads for 3 h at 4°C and washed thrice in a detergent-containing buffer (1 M Tris-HCl, pH 7.5, 5 M NaCl, 0.5 M EDTA, 0.1 M EGTA, and 1% Triton X-100). Immunoprecipitates were then boiled in SDS-containing loading buffer for 10 min prior to analysis by SDS-containing polyacrylamiden gel electrophoresis and Western blotting. Antibodies used for Western blotting were p97 (Invitrogen) and GFP (Clontech).

### Confocal microscopy and image analysis

One day prior to transfection, HC11 cells were seeded at 1 × 10^6^ cells in 6-well plates. Equal amounts of plasmids expressing GFP-SP WT, M57-58, or M59-61 were co-transfected with ER-mCherry using Lipofectamine 3000 (Thermo Fisher Scientific). At 24 h post-transfection, 6 × 10^5^ cells were seeded onto 22 × 22 1mm coverslips and, after another 24 h, fixed with 4% paraformaldehyde followed by permeabilization with 0.1% Triton-X 100. Cells were then washed thrice with PBS containing 0.1% Tween 20 (PBS-T). Blocking was performed with the same buffer containing 2% FBS. Fixed cells were incubated with 150 nM DAPI for 10 min followed by three washes with PBS-T. Finally, the coverslips were mounted on slides with VECTASHIELD mounting medium (Vector Laboratories) and sealed for imaging. The Zeiss LSM 710 Confocal and Elyra S.1 Structured Illumination Super Resolution Microscopes were used to capture Z-stack images of the samples. All slices in Z-stacks across samples were kept at 0.33 μm per slice interval. Lasers 405 nm, 488 nm, and 561 nm were used with the same power and gain across samples to capture images. Quantitation and image processing were performed using Zeiss software ZEN, Graphpad (Prism), and ImageJ. Briefly, Manders Colocalization Coefficient (MCC) (73) was used to quantify the percentage of GFP-SP pixels colocalized with ER-mCherry pixels in single slices. Mann-Whitney’s t tests were used to compare different samples in Graphpad software. Images were processed using ImageJ.

### Cloning and purification of GST-tagged SP proteins

GST-tagged wild-type or mutant SP expression constructs were generated by cloning the coding sequence into the *XhoI* and *NotI* restriction enzyme sites within the PGEX-6-GST vector. The constructs were transformed into BL21 cells, and a single colony was inoculated in 5 ml 2X YT media (16 g/L tryptone, 10 g/L yeast extract, and 5 g/L sodium chloride) containing ampicillin (100 μg/ml). After overnight incubation at 37°C with constant shaking at 200 rpm, the initial culture was inoculated into 500 ml of 2X YT media containing 100 μg/ml ampicillin and again grown at 37°C with constant shaking at 200 rpm to an optical density of ∼0.6. Induction of GST-SP proteins was performed by addition of 100 μM IPTG and overnight incubation at 16°C with constant shaking. The culture was then subjected to centrifugation at 7,000 rpm for 10 min at 4°C, and the pellet was lysed in PBS buffer containing 5 mM EDTA, 1X protease inhibitor cocktail (Thermo Fisher Scientific), 1 mM PMSF, and 1 mg/ml lysozyme. The lysate was centrifuged at 13,000 Xg for 30 min at 4°C, the cleared supernatant was collected and incubated with 100 μl of GST Sepharose beads (GE Healthcare) for 1-2 h at 4°C. The GST beads were washed four times with PBS. The bound protein was then eluted from the beads using PBS containing 10 mM reduced glutathione. The integrity and purity of the proteins were verified by SDS-polyacrylamide gel electrophoresis.

### Native gel electrophoresis

Self-association of GST-SP proteins was determined by native gel electrophoresis using Mini-PROTEAN TGX 4–20% precast polyacrylamide protein gels (Bio-Rad). Purified wild-type or mutant GST-SP (5 μg) were mixed with 2X native sample buffer (Laemmli buffer without SDS) (74) and subjected to electrophoresis using Tris-glycine running buffer at constant voltage (100V). Proteins were stained using Coomassie Blue. The results were repeated at least three times.

### Molecular modeling

Structural modeling of wild-type SP and mutants M57-58 and M59-61 was performed using the open-source ColabFold pipeline, which combines the multiple sequence alignment of MMseq2 with structure predictions by AlphaFold2 and RoseTTAFold (45, 46). Individual models were selected based on the average predicted local distance difference test (LDDT). The predicted structures were further visualized and analyzed using ChimeraX (47). The transmembrane helix was predicted using MEMSAT-SVM (48) and the coiled-coil was predicted using the CCHMM_PROF server using default settings (49).

### Statistical analyses and reproducibility

Either Mann-Whitney or Student’s t tests were performed as indicated. All experiments were performed at least twice with similar results.

## ACKNOWLEDGMENTS

We thank the Dudley laboratory for helpful discussions and comments on the manuscript. This work was supported by National Institutes of Health R01 grants CA167053 and AI131660.

